# Functional diversification of two Lon homologs enhances stress adaptation in *Pseudomonas aeruginosa*

**DOI:** 10.64898/2026.09.03.749167

**Authors:** Aswathy Kallazhi, Max Louski, Wissal Bakri, Efstathios Nikolaos Vlachos, Ute Römling, Kristina Jonas

## Abstract

The Lon protease is a highly conserved ATP-dependent protease that contributes to protein quality control and regulatory processes across all domains of life. The opportunistic pathogen *Pseudomonas aeruginosa*, along with other members of the Pseudomonadales, encodes AsrA (aminoglycoside-induced stress response ATP-dependent protease), a second LonA-type protease in addition to canonical Lon. Although AsrA is upregulated by aminoglycoside stress, its biochemical activity, substrate spectrum, and cellular functions have remained elusive. Here, we demonstrate that AsrA is a temperature- and antimicrobial stress-induced protease with specialized functions and partial redundancy with Lon. Quantitative proteomics revealed a distinct AsrA substrate profile, while comparative biochemical analyses showed that AsrA and Lon share a substantial number of substrates *in vitro* but differ in their degradation kinetics, indicating divergent substrate preferences. Among the proteins preferentially degraded by AsrA is the quorum-sensing anti-activator QslA, and we show that AsrA-dependent QslA degradation under tobramycin stress induces quorum-sensing gene expression. Together, our findings demonstrate how duplication of a conserved protease can generate specialized regulatory functions through differential expression and substrate preference, expanding the proteolytic network that enables bacterial adaptation to stress.

**Significance:** Proteolysis is central to bacterial stress adaptation, but how expansion of protease families generates new regulatory functions remains poorly understood. The opportunistic pathogen *Pseudomonas aeruginosa* encodes two homologous Lon proteases, Lon and AsrA. We show that AsrA has evolved distinct substrate preferences and stress-responsive functions while retaining substantial functional overlap with Lon. Notably, AsrA directly degrades the quorum-sensing anti-activator QslA during aminoglycoside stress, revealing a previously unrecognized connection between stress-induced proteolysis and quorum-sensing regulation. Our findings demonstrate how duplication and functional diversification of homologous proteases can expand bacterial regulatory capacity and promote adaptation to environmental stress. This work provides a framework for understanding how proteolytic network diversification contributes to bacterial adaptation.

## Introduction

Highly selective proteolysis is central to protein homeostasis and regulatory processes in all cell types. It is carried out by ATP-dependent proteases, many of which belong to the AAA+ (ATPases Associated with diverse cellular Activities) protein superfamily, that specifically recognize, unfold and degrade substrate proteins. In bacteria, proteases play key roles in remodelling the proteome in response to changing growth conditions, with a core set of proteases belonging to the Clp, Lon and FtsH protease families (1). While most bacteria rely on this set, some species have expanded this protease network with additional homologs of the canonical enzymes, often acquired through gene duplication in combination with vertical gene transfer or horizontal acquisition, allowing them to tailor their proteolytic systems to distinct cellular and environmental demands (2, 3).

One of the core proteolytic enzymes in bacteria is the Lon protease. By selectively degrading misfolded and damaged proteins, it plays a central role in maintaining protein homeostasis, especially under stress conditions (4). In addition, Lon fulfils important regulatory roles by targeting native proteins involved in a wide range of essential cellular processes, such as cell cycle progression, motility, stress responses, cell differentiation and pathogenicity(5–11). Lon homologs are also present in Archaea and in certain organelles of eukaryotic cells. For example, human mitochondrial Lon is essential for mitochondrial homeostasis and is critical for health (12).

The Lon enzyme assembles into a barrel-shaped hexameric complex composed of three functional domains, an N-terminal domain involved in substrate recognition, an AAA+ unfoldase domain as well as a C-terminal peptidase domain (4). Lon proteases are classified into three subfamilies, LonA, LonB and LonC, based on their domain organization, particularly the presence or absence of the N-terminal domain (13). While LonA contains all three domains and is widely found in both bacteria and eukaryotes, LonB and LonC lack the N-terminal domain, display subfamily-specific variations within their AAA+ domain, and are typically present in archaea or certain thermophilic bacteria, respectively (13).

A growing body of evidence indicates that many species encode Lon-like proteins or individual Lon domains in addition to their canonical Lon proteases, likely arising from ancient gene duplication events. In some cases, these Lon-like proteins exhibit domain architectures similar to the LonA, LonB or LonC proteases (14–18). In other instances, they consist of proteins resembling only one of the domains of Lon (19). A third group of proteins contain Lon-like domains fused to other functional domains, as exemplified by the bacterial RecA homolog RadA (20) and the CRISPR-associated Lon protease CalpL (21). With only few exceptions, the functions of Lon-like proteins, their evolutionary origins and their interplay with canonical Lon proteases, remain poorly studied.

The opportunistic pathogen *Pseudomonas aeruginosa* encodes a Lon-like protease (PA0779) in addition to its canonical Lon (LonA-type) protease (14). Owing to the strong upregulation of the corresponding gene upon exposure to the aminoglycoside antibiotic tobramycin, this protein was named AsrA (<u>a</u>minoglycoside-induced <u>s</u>tress response <u>A</u>TP-dependent protease) (14). Although a recent study suggests that AsrA may have certain redundant functions with Lon (22), its biochemical activity, substrate specificity and precise cellular roles remained to be defined.

Here, we functionally characterize the Lon homolog AsrA as the founding member of an evolutionary distinct subgroup of ATP-dependent LonA type proteases. Although AsrA shares the domain structure with Lon and also exhibits partly overlapping substrate specificity *in vitro*, it displays distinct substrate preferences. Most notably, we uncover a critical and specific role for AsrA in the antibiotic stress-dependent regulation of quorum sensing genes through the degradation of the anti-activator QslA. Together, our findings demonstrate how divergence of substrate preferences drives functional specialization of proteases and thereby allows expansion of a proteolytic network.

## Results

### AsrA proteins form a distinct subgroup of Lon proteases

*P. aeruginosa* encodes two LonA homologs, Lon and AsrA. While the *lon* gene (*PA1803*) resides within conserved genomic region alongside the protein quality control genes *tig*, *clpP* and *clpX*, the AsrA-encoding gene *PA0779* is located at a distinct genomic locus (**Fig. 1A**). Both genes have previously been reported to be under control of the heat shock sigma factor σ³² (22, 23). Hence, we assessed the expression of each protease under optimal conditions and following heat shock and tobramycin-induced aminoglycoside stress by immunoblotting using specific antisera, including an anti-AsrA antiserum raised for this study (**Fig. S1**). Lon was abundant under non-stress conditions and showed only a slight non-significant increase in response to heat shock and no obvious increase following tobramycin exposure (**Fig. 1B**). In contrast, AsrA was present at lower levels in the absence of stress but accumulated strongly in response to both proteotoxic stressors (**Fig. 1B**).

**Figure 1.**
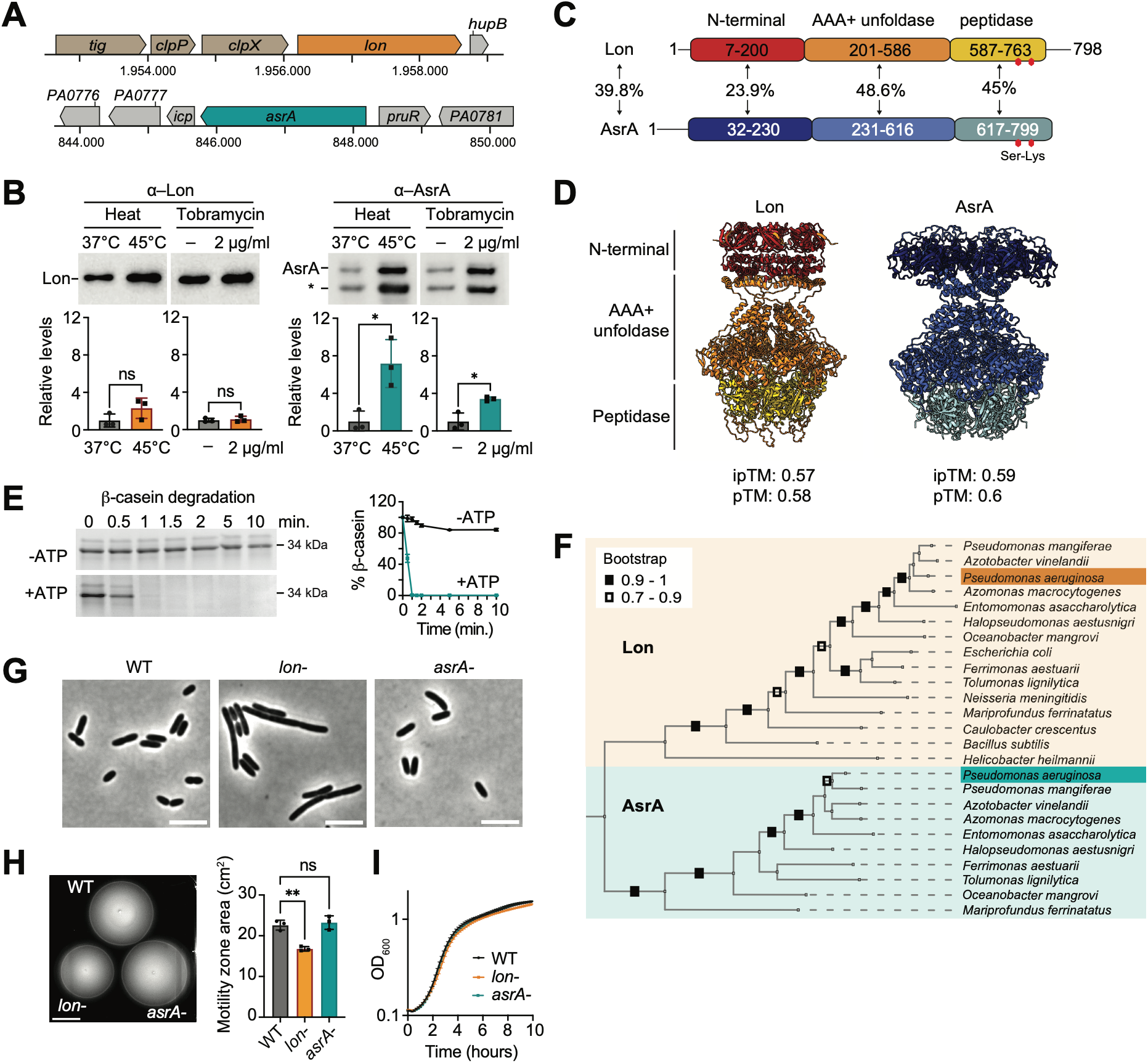
AsrA represents a distinct subgroup of the Lon protease family and is enzymatically active. **A)** Genomic context of *lon* (*PA1803*) and *asrA* (*PA0779*) in *P. aeruginosa* PAO1. **B)** Representative immunoblots showing Lon and AsrA in the *P. aeruginosa* PAO1 WT before and after heat stress and 2 μg/mL tobramycin treatment for 30 min using corresponding antibodies as indicated. Asterisk (*) depicts unspecific protein band. Loading control is shown in **Fig. S1B**. *Bottom:* Quantification showing the mean ± SD of three independent biological replicates. Statistical significance was determined using paired t-test for WT before and after stress for Lon and AsrA levels: Lon under heat stress, P=0.1654, ns; Lon under tobramycin treatment, P=0.3701, ns; AsrA under heat stress, P=0.0346, *; AsrA under tobramycin treatment, P=0.0246, * (ns – not significant). **C)** Domain structure of Lon and AsrA proteins according to UniProt. The percentage identity for the full-length proteins and individual domains is shown and based on Clustal Omega alignments. **D)** Side view of AlphaFold predicted 3D structure of Lon and AsrA hexamer (ipTM: interface predicted Template Modeling score; pTM: predicted Template Modeling score). **E)** *In vitro* degradation assay of β-casein in the presence (blue) and absence (black) of ATP and creatine kinase (CK). 0.2 μM of AsrA hexamer and 200 μg/mL of β-casein was used. Quantifications show the mean ± SD from three independent experiments. Band intensities of casein were normalized relative to corresponding AsrA levels for each time point. **F)** Phylogenetic tree constructed using maximum likelihood method in MEGA11 representing the homology between Lon and AsrA proteins of various classes of bacteria based on 1000 bootstrap replications. Bootstrap values of 0.7 or above are represented at the relevant nodes. **G)** Phase-contrast microscopy images of *lon-* strain and *asrA-* strains when grown at 37°C to exponential phase. Scale bar 5 μm. **H)** Swimming motility in soft agar after 48 h at room temperature of WT, *lon*-, and *asrA-* strains. Scale bar 2 cm. Quantification shows mean ± SD of motility area (cm^2^) from three independent biological replicates for each strain. Statistical significance was calculated using ordinary one-way ANOVA (Turkey’s multiple comparisons test) for comparison of pairs: WT vs *lon-*, P=0.0028, **; WT vs *asrA-*, P=0.8117, ns. **I)** Growth curve of WT, *lon-*, and *asrA-* strains grown at 37°C. The means ± SD of three independent biological replicates are plotted.

Comparison of the amino acid sequence and AlphaFold-predicted structure of AsrA with *P. aeruginosa* Lon revealed that, although AsrA and Lon share only 39.8% sequence identity (**Fig. 1C**, **Fig. S2**), they adopt highly similar structures, both as predicted monomers and as assembled hexameric complexes (**Fig. 1D**, **Fig. S3**). The sequence divergence is particularly pronounced in the N-terminal domain, which exhibits only 23.9% sequence identity (**Fig. 1C**), and displays greater structural variation than the remainder of the protein (**Fig. 1D**). AsrA retains intact Walker A and B motifs within the AAA+ domain important for ATP binding and hydrolysis as well as the serine-lysine catalytic dyad in the peptidase domain (**Fig. 1C**, **Fig. S2**). However, its proteolytic activity had not been experimentally demonstrated. To address this, we purified AsrA and assessed its ability to degrade the model substrate β-casein. Rapid degradation was observed in the presence of ATP, but not in its absence, confirming that AsrA is an ATP-dependent protease (**Fig. 1E**).

Retrieving the closest AsrA homologs revealed that AsrA is particularly widespread within the order *Pseudomonadales*, but is also present in the orders *Alteromonadales*, *Aeromonadales* and *Oceanospirillales* as well as in some species outside the Gammaproteobacteria, such as the zetaproteobacterium *Mariprofundus ferrinatatus*. Phylogenetic analyses of AsrA and Lon proteases from selected bacterial classes and orders showed that the two proteins form two distinct clades (**Fig. 1F**). This separation into the Lon and AsrA clusters remained evident when including the more distantly related Lon homologs from *Bacillus subtilis* and *Caulobacter crescentus*, indicating that Lon and AsrA represent ancient subgroups within the Lon protease family. This divergence suggests that the two proteases diverged early in evolution and may have evolved to fulfil distinct cellular functions. Consistent with this idea, transposon disruption mutants of *asrA* and *lon* display different phenotypes under non-stress conditions: while the *lon-* mutant exhibits impaired motility, slightly reduced growth and severe cell filamentation due to accumulation of the cell division inhibitor SulA (9) (**Fig. 1G-I**), we did not detect these defects in the *asrA-* mutant (**Fig. 1G-I**).

### Quantitative proteomics reveals novel substrates of AsrA

While recent proteomics studies have identified numerous protein substrates of Lon in several species (8, 9, 24, 25), direct target proteins of AsrA remained unknown. To uncover specific AsrA substrates as well as proteins indirectly affected by its activity, we used mass spectrometry-based quantitative proteomics to analyze changes in protein steady-state levels and stabilities in strains either overexpressing or lacking AsrA. Specifically, we searched for proteins whose abundance and stability decreased upon overexpression of *asrA* and that showed an increased steady-state level in the *asrA-* mutant by three different approaches. To identify proteins with reduced abundance due to *asrA* overexpression, we compared the proteome of a strain overexpressing *asrA* for 90 min with that of a vector control (VC) strain harbouring an empty vector. To assess the effect of AsrA on protein stability, we inhibited protein synthesis with spectinomycin and monitored protein levels after 60 min in the vector control and *asrA* overexpression strains. Conversely, in order to identify proteins accumulating in the absence of *asrA*, we compared the proteomes of an *asrA-* mutant and the corresponding wild type strain following 90 min of tobramycin stress. Protein quantification across all conditions using TMT (Tandem Mass Tag) isobaric labelling resulted in the detection of in total 2021 proteins. Of these, 926 proteins showed reduced steady-state levels in *asrA-* overexpressing cells compared to the vector control strain, and 992 proteins showed increased levels in the *asrA-* mutant under tobramycin stress (**Fig. 2A, Dataset S1**). Additionally, 614 proteins were found to be unstable in the *asrA* overexpression strain. To identify potential direct AsrA substrates, we focused on proteins meeting all three criteria, resulting in a shortlist of 124 proteins (**Fig. 2B, Dataset S1**). Functional classifications of these proteins revealed a broad spectrum of categories, including many metabolic proteins as well as transporters and transcriptional regulators (**Fig. 2C**). Comparison of the 124 putative AsrA substrates with previously reported Lon substrate candidates identified through comparable quantitative proteomics-based approaches (9) revealed surprisingly little overlap (**Fig. 2D**). Only eight proteins were common to both datasets, and among these, FliG, FliS, and SpeH have been experimentally validated as *bona fide* Lon substrates (9).

**Figure 2.**
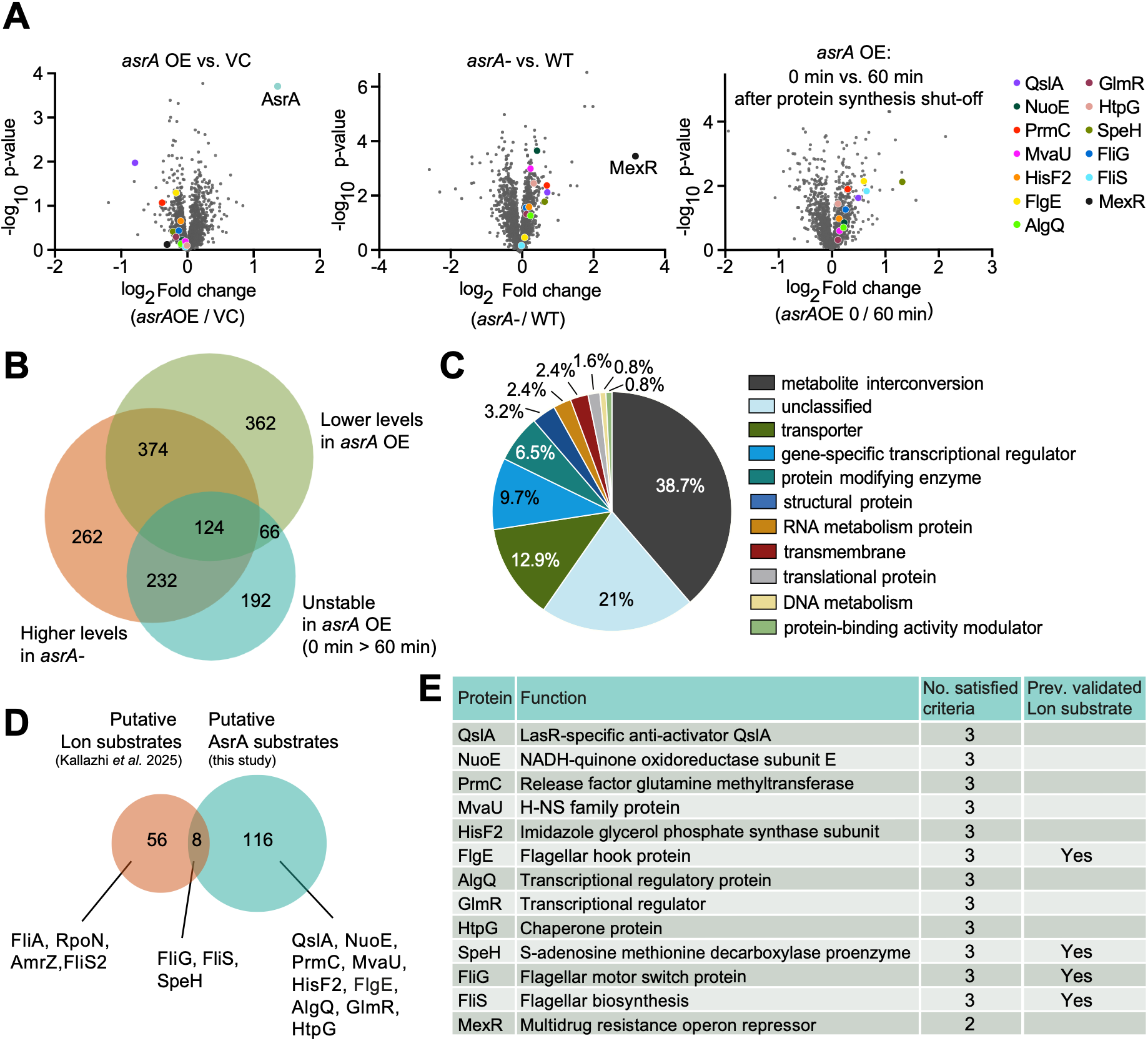
A quantitative proteomics approach reveals novel AsrA substrates. **A)** Volcano plots showing proteome-wide effects in response to *asrA* overexpression (*asrA*OE) in comparison to the vector control (VC) after 90 min of *asrA* induction (*left*), in the *asrA*-strain compared to the wild type (WT) after 90 min of tobramycin treatment (*middle*) and 60 min after spectinomycin-induced synthesis shut-down in the *asrA*OE strain in comparison to the 0 min time point (*right*). Selected proteins investigated in this study are marked in all the plots. p-values were calculated using a two-tailed t-test based on three independent biological replicates. **B)** Venn diagram showing filtering of substrate candidates by applying three different criteria. Criterion I, green circle — proteins showing a lower steady-state level in *asrA*OE compared to the VC. Criterion II, orange circle — proteins showing higher levels after 90 min of tobramycin addition in *asrA-* compared to the WT. Criterion III, blue circle — proteins that are unstable in *asrA*OE, i.e., showing a downregulation between 0 to 60 min. **C)** Pie chart of the functional classification analysis of the 124 proteins satisfying all the criteria in Fig. 2B. **D)** Venn diagram showing the overlap of the 64 putative Lon substrates reported by Kallazhi *et al.*(9) and the 124 putative AsrA substrates identified in this study. **E)** Table of selected putative AsrA substrates with their primary annotated function and number of satisfied criteria. The right column indicates whether each protein was previously (prev.) validated as a Lon substrate by Kallazhi *et al.* (9).

### Lon and AsrA have largely overlapping substrate specificity but divergent substrate preferences

Based on the results of the proteomics experiment, we selected a group of 13 putative AsrA substrates for further analysis (**Fig. 2E**). This group included 12 candidates that satisfied all three criteria to varying extents and included the four previously validated Lon substrates FlgE, FliG, FliS and SpeH (9). Additionally, we included MexR, a repressor of the MexAB-OprM multidrug efflux pump and thus critical determinant of antibiotic resistance in *P. aeruginosa* (26), which was the most strongly upregulated protein in the *asrA-* strain and was also downregulated upon *asrA* overexpression (**Fig. 2A**), but did not meet the criterium of being unstable in the *asrA* overexpression strain.

We purified the substrate candidates and monitored their degradation by AsrA using *in vitro* degradation assays. While we did not detect ATP-dependent degradation of MexR, GlmR, HtpG and AlgQ (**Fig. 3A**), the other nine substrates (QslA, NuoE, PrmC, HisF2, SpeH, FliG, MvaU, FliS and FlgE) showed robust degradation by AsrA in the presence of ATP (**Fig. 3B**) but not in its absence (**Fig. S4A**), confirming them as direct AsrA substrates. Some of these AsrA substrates, like NuoE and PrmC, showed notably fast degradation and were completely cleared within 40 minutes, while others (e.g. HisF2 or SpeH) showed more modest degradation. Given the structural similarity between Lon and AsrA (**Fig. 1D**), and the fact that four of the proteins have previously been shown to be Lon substrates, we compared the degradation of all nine proteins by AsrA and Lon side-by-side. Interestingly, all nine validated AsrA substrate proteins were also degraded by Lon (**Fig. 3B**), whereas the four proteins that were not degraded by AsrA (MexR, GlmR, HtpG, AlgQ) were likewise resistant to Lon-mediated degradation (**Fig. S4B**). Together, these *in vitro* results indicate a high degree of overlap in substrate specificity between AsrA and Lon. Importantly, however, quantification of degradation kinetics demonstrated substrate-specific kinetic differences between the two proteases (**Fig. 3C**). NuoE, PrmC, FliG, FliS, HisF2 and FlgE were degraded substantially faster by AsrA than by Lon, whereas MvaU and SpeH were degraded more efficiently by Lon. In contrast, QslA, was degraded with similar kinetics by both proteases.

**Figure 3.**
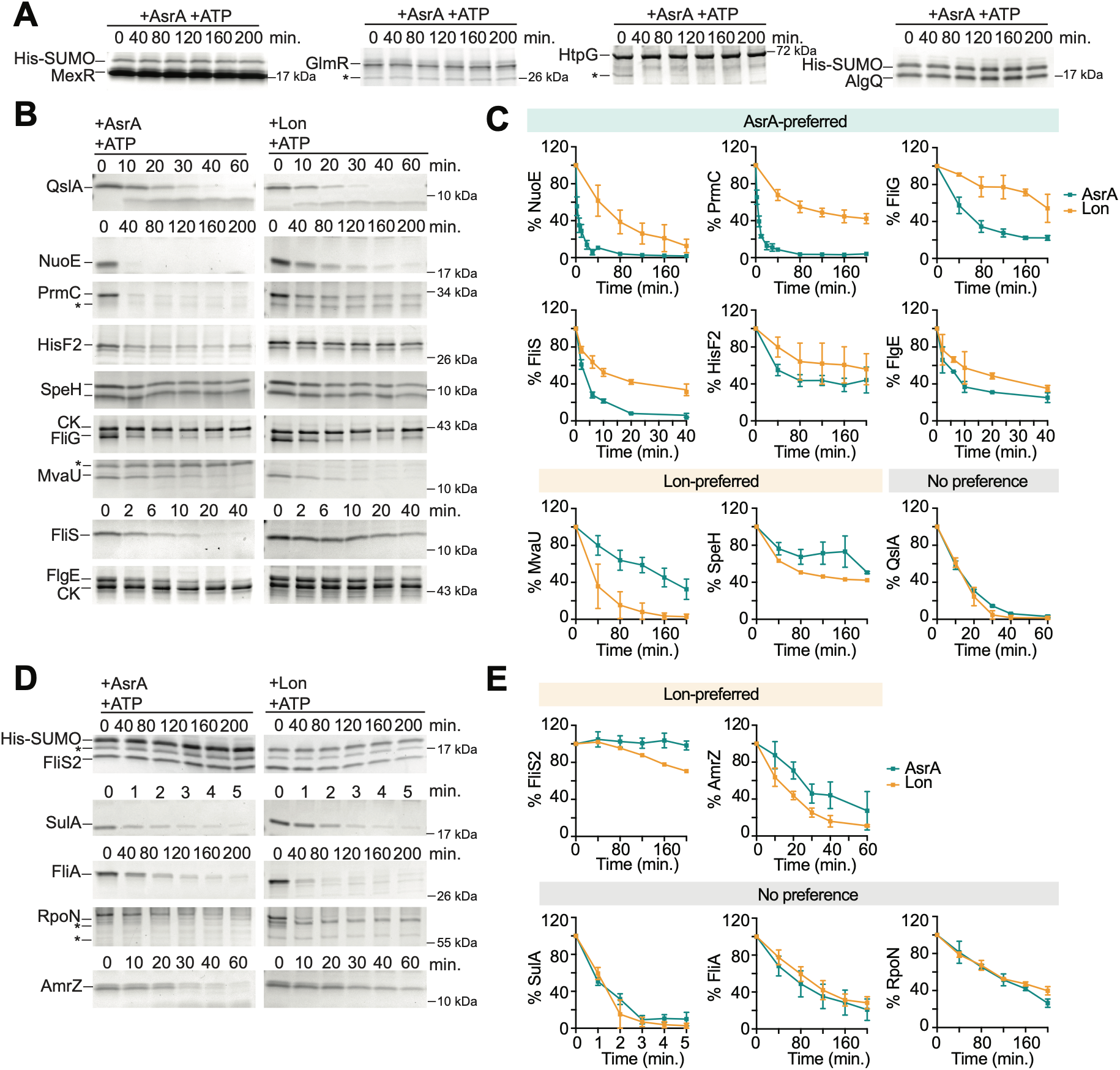
Lon and AsrA share a large set of substrates *in vitro*, but degrade them with distinct kinetics. **A)** *In vitro* degradation assays of MexR, GlmR, HtpG, and AlgQ in the presence of AsrA, ATP, and creatine kinase (CK). 0.4 μM of AsrA hexamer and 4 μM of MexR. 0.2 μM of AsrA hexamer and 2 μM of GlmR, HtpG, and 3 μM of AlgQ were used. One representative SDS-PAGE gel of each assay is shown, with molecular weight of closest ladder band indicated. Asterisks (*) depict unspecific protein bands. **B)** *In vitro* degradation assays of QslA, NuoE, PrmC, HisF2, SpeH, FliG, MvaU, FliS, and FlgE in the presence of AsrA or Lon with ATP and creatine kinase. 0.2 μM of AsrA or Lon hexamer and 2 μM of QslA, 8 μM of NuoE, 2 μM of PrmC, 2 μM of HisF2, 4 μM of SpeH, 1 μM of FliG, 2 μM of MvaU, 1.5 μM of FliS, and 0.75 μM of FlgE were used. 0.4 μM of AsrA hexamer and 4 μM of QslA and 4 μM of HisF2 was used. One representative SDS-PAGE gel of each assay is shown. **C)** Quantifications of *in vitro* degradation assays shown in Fig. 3B. The relative substrate levels are normalized to corresponding Lon/AsrA levels at each time point. Quantifications show the mean ± SD of at least two independent replicates and are sorted according to proteases preference. **D)** *In vitro* degradation assays of the previously validated Lon substrates FliS2, SulA, FliA, RpoN, and AmrZ in the presence of AsrA or Lon, ATP, and creatine kinase. 0.2 μM of AsrA or Lon hexamer and 1.5 μM FliS2, FliA, RpoN, and 2 μM of SulA. One representative SDS-PAGE gel of each assay is shown. See **Fig. S4C** for corresponding *in vitro* degradation assays in the absence of ATP. **E)** Quantification of *in vitro* degradation assays shown in Fig. 3D. Quantifications were done and represented as in Fig. 3C.

Prompted by these results, we also tested if AsrA can degrade FliS2, SulA, FliA, RpoN and AmrZ, five additional previously reported Lon substrates (9), which however did not meet all three criteria for being classified as a putative AsrA substrate in the proteomics experiment (**Fig. 2D**, **Fig. S5**). While FliS2 was not degraded by AsrA over 200 min, the other four proteins were degraded by AsrA in an ATP-dependent manner (**Fig. 3D**, **Fig. S4C**). Of these proteins, SulA, FliA and RpoN were cleared by AsrA and Lon with similar kinetics, while AmrZ was degraded by AsrA more slowly than by Lon (**Fig. 3E**).

Together, our biochemical analysis revealed 13 proteins as native substrates of AsrA. Our data indicate a large overlap in substrate specificity between AsrA and Lon. However, differences in *in vitro* degradation kinetics of some substrates point to distinct substrate preferences. These substrate preferences may reflect specialized roles of Lon and AsrA *in vivo*, potentially modulated by accessory adaptors, regulatory proteins or small molecules.

### AsrA upregulates quorum sensing genes by degrading the anti-activator QslA

One of the newly identified Lon and AsrA substrates was QslA (PA1244) that was rapidly degraded by both proteases *in vitro*. QslA is the anti-activator of LasR, a master regulator of quorum sensing signalling in *P. aeruginosa* (27), which senses the 3O-C12-homoserine lactone autoinducer to activate gene expression. By directly binding LasR, QslA prevents LasR dimerization and subsequent DNA binding (28), thereby repressing a large set of quorum sensing genes, including the *pqs* genes responsible for PQS (*Pseudomonas* Quinolone Signal) biosynthesis (29) (**Fig. 4A**). Based on our proteomics data, QslA was among the most promising AsrA substrate candidates that was clearly upregulated in *asrA-* cells and, conversely, downregulated following *asrA* overexpression (**Fig. 4B**), but was not shortlisted as a potential Lon substrate according to our previous study (9).

**Figure 4.**
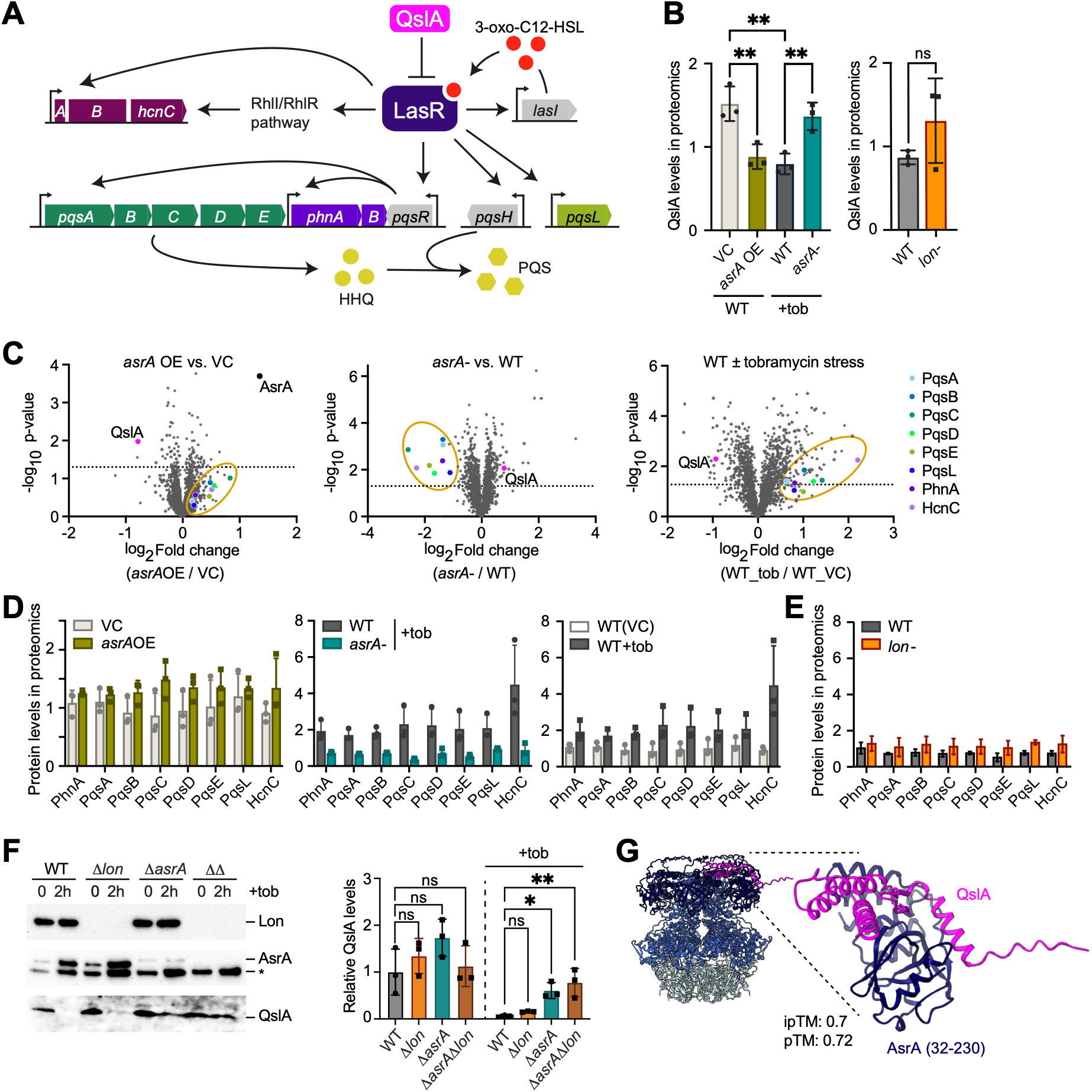
AsrA regulates the PQS quorum sensing pathway via degradation of the anti-activator QslA. **A)** Simplified schematic illustration of the LasR-controlled PQS quorum-sensing pathway in *P. aeruginosa*. **B)** Left: QslA levels in the proteomics data in response to *asrA* overexpression (*asrA*OE) compared to the vector control (VC) and in the *asrA*-strain compared to the WT in the presence of tobramycin (+tob). The mean ± SD of at least three independent biological replicates are plotted. Statistical significance was calculated using one-way ANOVA (Tukey’s multiple comparisons test) for comparison of pairs: VC vs *asrA*OE P_adj_ = 0.0044, **; WT vs *asrA*-P_adj_ = 0.0083, **; VC vs WT P_adj_ = 0.0020, **. *Right*: QslA levels in the proteomics data from Kallazhi *et al.* (9) in the *lon-* strain in comparison to the WT. The mean ± SD of at least three independent biological replicates. Statistical significance was calculated using an unpaired t-test for comparison of WT vs *lon-*, P=0.2129, ns (ns: not significant). **C)** Volcano plots of the proteomics data showing the change of QslA and other quorum sensing proteins in response to *asrA* overexpression versus VC (*left*), *asrA-* versus WT (+tobramycin) (*middle*) and in response to tobramycin stress in WT (*right*). In the condition (–tobramycin), the WT contained the empty vector control (VC). The dotted line indicates −log_10_ p-value of 1.3 (corresponding to a p-value of 0.05). **D)** Protein levels of PhnA, PqsA, PqsB, PqsC, PqsD, PqsE, PqsL and HcnC as determined in the proteomics experiment for the same conditions as shown in Fig. 4C (*right*). **E)** Protein levels of PhnA, PqsA, PqsB, PqsC, PqsD, PqsE, PqsL and HcnC as determined by a previous proteomics experiment(9) in WT and *lon-*. **F)** Representative immunoblots (*left*) showing Lon, AsrA and QslA levels in the WT, Δ*lon*, Δ*asrA*, Δ*asrA* Δ*lon* (ΔΔ) strains before and after 2 μg/mL tobramycin treatment for 2 hours using corresponding antibodies as indicated. Asterisk (*) depicts unspecific protein band. Loading control is shown in **Fig. S6**. Quantification (*right*) sowing mean ± SD based on three independent biological replicates. Statistical significance was calculated using ordinary one-way ANOVA (Turkey’s multiple comparisons test) for comparison of pairs: WT vs Δ*lon* before tobramycin treatment, P_adj_ =0.7619, ns; WT vs Δ*asrA* before tobramycin treatment, P_adj_ =0.2307, ns; WT vs Δ*asrA* Δ*lon* before tobramycin treatment, P_adj_ =0.9826, ns; WT vs Δ*lon* after tobramycin treatment, P_adj_ =0.9391, ns; WT vs Δ*asrA* after tobramycin treatment, P_adj_ =0.02307, *; WT vs Δ*asrA* Δ*lon* after tobramycin treatment, P_adj_ =0.005, **. **G)** Structural model of the Alphafold3 predicted QslA–AsrA complex. Structural superposition of the predicted QslA–AsrA NTD binary complex onto the full AsrA homohexamer, illustrating the spatial orientation of QslA relative to the complete ring assembly. Close-up view shows the predicted direct interaction between QslA (magenta) and a single AsrA NTD subunit (dark blue).

Given QslA’s role as a negative regulator of LasR, we asked whether AsrA-dependent QslA degradation affects the LasR regulon. Indeed, the most strongly downregulated proteins in the *asrA-* strain were PqsABCDE as well as PqsL, PhnA and HcnC (**Fig. 4C**, **D**), which are all known to be positively controlled by LasR (30) (**Fig. 4A**). Conversely, the same set of genes was upregulated in *asrA* overexpressing cells, in which QslA levels are reduced. Because the *asrA*-proteomics data were collected under tobramycin stress, we wondered whether this condition makes AsrA-dependent degradation of QslA particularly important. Supporting this idea, the *pqs* genes were upregulated in wild-type cells in response to tobramycin, consistent with previous observations (14) (**Fig. 4C**, **D**), suggesting that AsrA-mediated QslA degradation contributes to the induction of quorum sensing genes under stress.

Since QslA was also degraded by Lon *in vitro*, we investigated how quorum sensing genes were affected by Lon by analyzing a previous proteomics dataset which compared protein levels between WT and *lon*-strain (9). While QslA levels were mildly but non-significantly upregulated in the *lon*-mutant (**Fig. 4B**), none of the PQS proteins were significantly reduced in abundance; in fact, the abundance of PQS proteins was instead slightly elevated in *lon*-cells (**Fig. 4E**). Because the previous *lon*-proteomics data were obtained in the absence of tobramycin, we generated an anti-QslA antiserum to investigate the effects of AsrA and Lon on QslA levels more thoroughly *in vivo* by immunoblotting. More specifically, we analyzed QslA abundance in recently published clean deletion mutants lacking *lon*, *asrA*, or both genes (Δ*asrA* Δ*lon*) (22), and compared them with the wild type before and after tobramycin exposure. While QslA was almost completely eliminated in the wild type and the Δ*lon* mutant following tobramycin treatment, it remained clearly detectable in the Δ*asrA* mutant and was maintained in the Δ*asrA* Δ*lon* double mutant at levels close to those observed under non-stress conditions (**Fig. 4F**, **Fig. S6**). These findings are consistent with the proteomics data and suggest that despite similar degradation kinetics of QslA by both proteases *in vitro*, AsrA is the primary protease responsible for QslA degradation *in vivo* upon tobramycin exposure, with Lon contributing primarily in the absence of AsrA.

Prompted by our findings, we used AlphaFold3 to assess whether it predicts a physical interaction between QslA and the N-terminal domains (NTDs) of AsrA and Lon. We obtained a high-confidence model for the AsrA NTD–QslA complex (ipTM score 0.7), in which the three predicted α-helices of QslA contact the surface of the globular region of AsrA’s NTD (**Fig. 4G**). In contrast, no high-confidence interaction was predicted for QslA with the NTD of Lon, suggesting a potentially more favourable interaction between QslA and AsrA, which is consistent with our *in vivo* observations.

Taken together, our results indicate that AsrA positively regulates the *pqs* quorum sensing system by degrading the anti-activator QslA, thereby promoting quorum sensing signalling pathways. Moreover, our data suggest that this regulatory role is particularly important for the previously observed induction of the PQS pathway under aminoglycoside stress (14). While Lon can also degrade QslA *in vitro* and may contribute to its turnover *in vivo*, our data indicate that AsrA is the preferred protease for QslA and is primarily responsible for its clearance upon exposure to tobramycin-induced stress.

### Redundant roles of Lon and AsrA in heat and tobramycin-induced stress survival

Given the well-recognized role of Lon proteases in protein quality control during stress conditions (4) and the large overlap in substrate specificity between AsrA and Lon (**Fig. 3**), we sought to assess their contributions to viability under non-stress conditions as well as proteotoxic stress survival. Under optimal growth conditions, only the Δ*lon* single mutant and the Δ*asrA* Δ*lon* double mutant showed mildly reduced colony formation, likely due to cell filamentation resulting from impaired Lon-dependent SulA degradation (9). Upon exposure to heat stress or sublethal tobramycin concentrations, neither the Δ*lon* nor the Δ*asrA* single mutants exhibited significant changes in survival. In stark contrast, the double mutant showed severely impaired survival under both heat and tobramycin stress (**Fig. 5A**). Although less pronounced, the double mutant also displayed a clear growth defect when grown in liquid medium in the presence of tobramycin compared with both the single mutants and the wild type (**Fig. 5B**). These results indicate that AsrA and Lon play partially redundant roles in protecting cells from heat- and tobramycin-induced stress. This functional overlap likely stems from the coordinated degradation of some of the substrates described here, additional substrates that remain to be identified, and/or the clearance of damaged or misfolded proteins.

**Figure 5.**
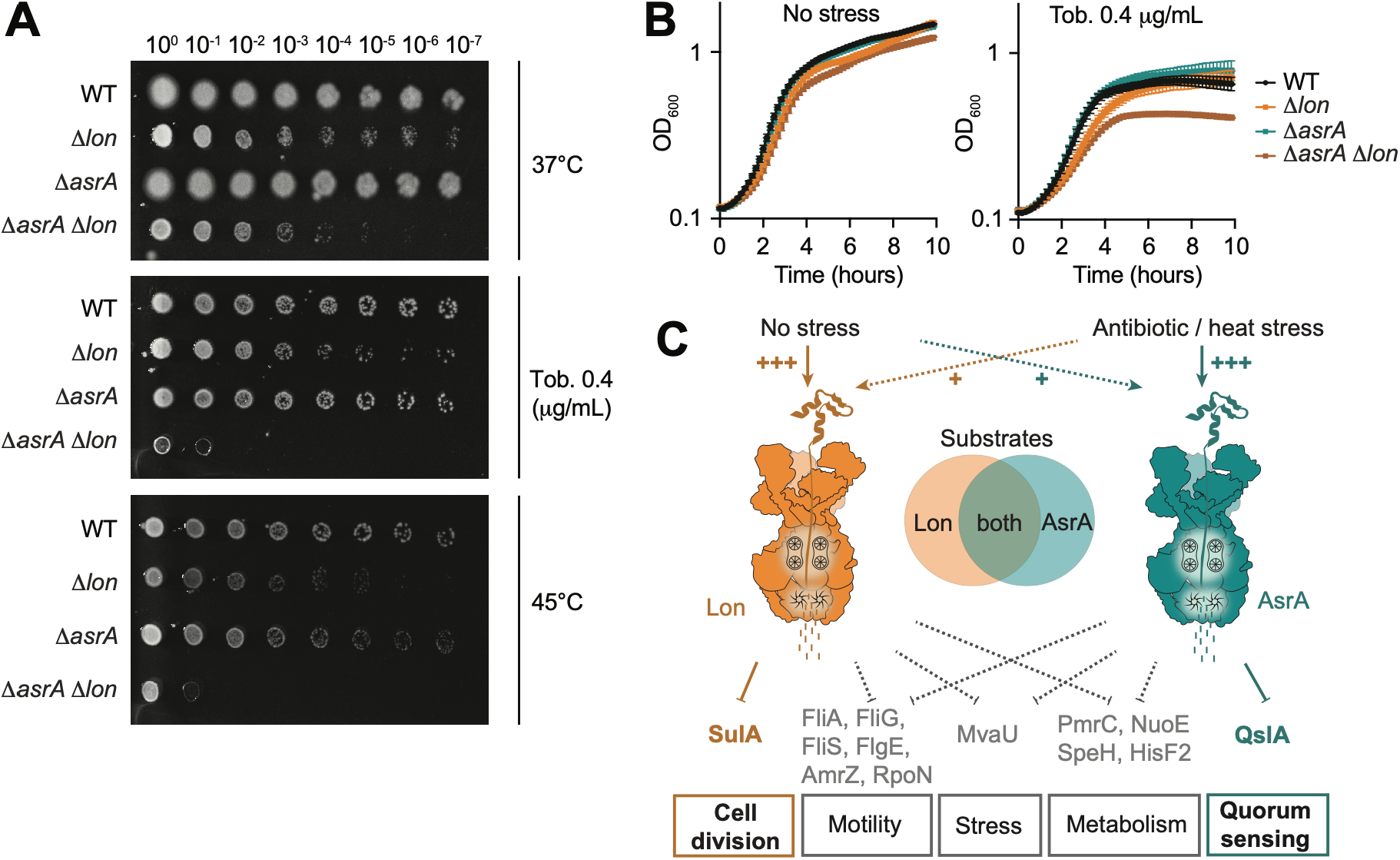
Lon and AsrA play redundant roles in survival during heat and tobramycin stress. **A)** Spot dilution assays of *P. aeruginosa* PAO1 WT, Δ*lon*, Δ*asrA*, Δ*asrA* Δ*lon* incubated overnight at 37°C on LB agar under no stress, on LB agar supplemented with tobramycin, or at 45°C on LB agar. Dilutions are spotted up to 10^-7^ dilution. **B)** Growth curves of the WT, Δ*lon*, Δ*asrA*, Δ*asrA* Δ*lon* grown at 37°C in LB without stress and under tobramycin 0.4 μg/mL. Means ± SD of at least three independent biological replicates are plotted. **C)** Schematic model showing the interplay and functional specialization of the Lon and AsrA proteases. The two proteases are differentially regulated in response to aminoglycoside and heat-induced proteotoxic stress. They degrade a shared set of substrates, involved in motility, stress responses, and metabolism. In addition, their distinct substrate preferences confer pathway-specific functions. Cell division is primarily regulated by Lon through degradation of SulA, whereas AsrA specifically regulates quorum sensing by degrading QslA.

## Discussion

Here, we establish the Lon-like protease AsrA as a distinct subgroup within the Lon protease family, present in members of the *Pseudomonadales* and related orders of the Gammaproteobacteria. Although AsrA and canonical Lon proteases share the same domain architecture and exhibit largely conserved overall structures, their sequence identity is relatively low, particularly within the N-terminal domain (NTD), which is involved in substrate recognition. Together with our phylogenetic analysis, this suggests that AsrA may have originated relatively early in evolution, likely through gene duplication followed by divergent evolution and subsequent speciation, potentially combined with horizontal gene transfer events.

By focusing on AsrA from *P. aeruginosa*, we characterized this protease in detail, identified and validated its first native substrates, and examined its functional relationship with Lon. Although AsrA and Lon belong to distinct subgroups of the LonA protease family, our *in vitro* analyses revealed substantial overlap in their substrate repertoires (**Fig. 5C**). Furthermore, simultaneous deletion of both proteases resulted in a synthetic survival defect under stress conditions, indicating that they cooperate to support survival during heat and antibiotic stress. Despite this apparent functional redundancy, our data also indicate specialization. AsrA appears to be dedicated to stress adaptation as its expression is strongly induced by antibiotic and heat stress. In contrast, Lon is abundant even under non-stress conditions consistent with a constitutive housekeeping role. Moreover, our biochemical analyses revealed distinct substrate degradation kinetics for AsrA and Lon, while our *in vivo* proteomic and phenotypic data further demonstrated that the two proteases differentially shape the cellular proteome and fulfill specialized physiological functions (**Fig. 5C**).

One example of a differentially affected substrate is the cell division inhibitor SulA. Although both proteases efficiently degrade SulA *in vitro*, specifically the loss of Lon results in its accumulation *in vivo*, leading to impaired cell division and consequent defects in growth and motility (9). In contrast, AsrA lacking cells do not exhibit cell division defects and our proteomics data indicate that they do not accumulate SulA. The apparent discrepancy between the *in vitro* and *in vivo* data suggests that substrate recognition and degradation may be influenced by additional cellular factors that modulate protease specificity or accessibility under physiological conditions. In several ATP-dependent protease systems, substrate selectivity is controlled by adaptor proteins that facilitate substrate recognition and delivery (31). In the case of Lon, however, only a limited number of such regulators have been identified and characterized, including HspQ in enteric bacteria (32), LarA in *Caulobacter crescentus* (33), and SmiA in *Bacillus subtilis* (10). In *P. aeruginosa*, SadB has recently been proposed to influence Lon-dependent degradation of AmrZ (34), although the precise underlying mechanism remains unclear. Our findings raise the possibility that analogous regulatory factors exist for AsrA and Lon and may differentially direct substrate selection, thereby contributing to their overlapping yet distinct cellular functions. The fact that the N-termini of these proteases, which are often the target of these regulatory factors, also share the least amount of conservation adds further strength to this hypothesis.

Another substrate exhibiting differential regulation by AsrA and Lon is QslA, which operates at the top of the QS hierarchy (27). Although both proteases efficiently degrade QslA *in vitro*, AsrA plays the predominant role in its turnover *in vivo*. In fact, the result that *pqs* genes were the most strongly downregulated in the *asrA-* mutant under tobramycin stress suggests that a key function of AsrA under this condition is to promote quorum sensing (QS) through the degradation of QslA. *P. aeruginosa* relies on a highly sophisticated QS network for environmental adaptation and pathogenicity, which is considered an attractive antibiotic drug target (35). Our finding of AsrA-dependent QslA regulation adds an additional layer of control to this system and expands our understanding of QS regulation in this organism. While our data suggest that Lon contributes only modestly to QslA regulation, previous work showed that LasI accumulates and is stabilized in a *lon* disruption mutant (36). Although direct *in vitro* degradation of LasI by Lon has not been demonstrated, these observations indicate that Lon also influences the QS network, albeit at a different regulatory level than AsrA.

Beyond SulA and QslA, our study identifies a broader set of functionally diverse AsrA and Lon substrates, including proteins involved in metabolism and gene regulation, such as NuoE, PrmC, HisF2, and MvaU (**Fig. 5C**). Given their diverse cellular functions, the coordinated degradation of these substrates likely contributes to the distinct physiological roles of AsrA and Lon and may also account for the increased stress sensitivity of the Δ*lon* Δ*asrA* double mutant. This may be particularly relevant for NuoE, a subunit of NADH:ubiquinone oxidoreductase (Complex I) in the respiratory chain, whose function, together with other respiratory components, has been associated with aminoglycoside tolerance through altered membrane potential (14, 37). Future studies will be required to elucidate how selective degradation of these proteins contributes to stress adaptation.

In conclusion, our work on AsrA illustrates how bacterial protease networks can expand through the emergence of homologous enzymes that retain partially overlapping yet functionally specialized roles. While AsrA represents one example of a Lon-like protease, bacterial genomes encode a wide variety of Lon-related proteins (14–21), and recent studies have begun to uncover the diverse functions of this expanding protein family. Similar patterns are observed for other major chaperone and protease systems, which frequently include additional homologs alongside their canonical counterparts (2, 3). Notable examples are the horizontally acquired copies of ClpG and FtsH found in *P. aeruginosa* Clone C, which perform complementary or additive functions relative to the canonical proteins and are thought to have contributed substantially to the clonal expansion and ecological success of this lineage (38, 39). These observations suggest that the expansion of protease and chaperone networks is a recurring evolutionary strategy that enhances bacterial adaptability.

## Materials and Methods

### Bacterial strains and plasmids

All bacterial strains used in this study are listed in Supplementary Table S1.

### Growth conditions

*P. aeruginosa* PAO1 and mutant strains were routinely grown in LB medium at 37°C while shaking at 200 rpm unless otherwise indicated. For all experiments involving induction of *asrA*, 0.2% L-arabinose was added to both the overexpression strain as well as the control strain with the empty vector pJN105. When needed, the medium was supplemented with either gentamicin (30 µg/ml) or tetracycline (50 µg/ml).

The salt-inducible BL21-SI/pCodonPlus *E. coli* strain for protein expression was grown at 30°C using LBON/2xYTON no salt medium. *E. coli* DH5α strain used for cloning was grown at 37°C in LB medium. When necessary, media was supplemented with antibiotics at following concentrations in liquid/solid (µg/ml): gentamicin (15/20), kanamycin (30/50) and chloramphenicol (20/40).

### Plasmid construction

All plasmids used in this study are listed in Supplementary Table S2 and primers in Supplementary Table S3. The cloning procedures for plasmids generated in this study are described in the Supplementary Methods.

### Sequence alignment of Lon and AsrA orthologs

Alignment of Lon and AsrA protein sequences was generated using EMBL-EBI Multiple Sequence alignment tool Clustal Omega (40) and visualized using Jalview 2.11.5.1 (41). Protein sequences were retrieved from ncbi.nlm.nih.gov.

### Protein structure prediction

Lon and AsrA 3D structure predictions were generated using the AlphaFold3 web server (https://alphafoldserver.com). The AsrA NTD–QslA complex was predicted using a local implementation of AlphaFold3 on an HPC cluster with model parameters provided by Google DeepMind (42). The resulting structural models were visualized and analyzed using UCSF ChimeraX (v1.5) (43).

### Phylogenetic analysis

The experimental clustered non-redundant database from Protein BLAST was used to collect 500 closest homologs of AsrA (RefSeq: NP_249470.1) and Lon (RefSeq: NP_250494.1). At least one sequence of an AsrA homolog from each of the largest clades obtained were selected. The phylogenetic tree was constructed using a MUSCLE multiple sequence alignment (44) and the maximum likelihood method using the software MEGA11 (45). The robustness was tested using a bootstrap analysis of 1000 replications. The obtained tree was visualized and edited using the online tool Interactive Tree Of Life (https://itol.embl.de/) (46).

### Sample collection and preparation for proteomics experiments

For the proteomics experiment, samples were collected and prepared as follows. For the samples involving overexpression of *asrA,* cultures of *P. aeruginosa* PAO1 cells containing either the empty vector pJN105 or pJN105:*asrA* were grown in triplicates of 200 mL LB cultures containing gentamicin for plasmid maintenance, into exponential phase cultures by back diluting 3–4 times. After collecting 1 mL pre-induction samples for immunoblot analysis, 0.2% arabinose was added to both cultures and grown at 37°C for 90 min. 1 mg/ml spectinomycin was then added to both the cultures and 50 mL of the culture was immediately retrieved into pre-cooled 50 mL tubes, spun down in a pre-cooled centrifuge and the supernatant discarded (0 min samples). The pellet was frozen using liquid nitrogen and stored at –20°C. OD_600_ values were measured throughout the experiment at hourly intervals to ensure comparable growth across all strains and replicates. The sample collection was repeated at 60 min after addition of spectinomycin. Three replicates each of 0 and 60 min samples were collected.

For the same proteomics experiment, in order to sample WT and *asrA*-strains under tobramycin stress conditions, *P. aeruginosa* strain PAO1 WT and the transposon mutant *asrA-* were grown in triplicates. After back-diluting 3–4 times, the exponential phase cultures were treated with 2 μg/mL tobramycin and samples were collected at 90 min as described before. Three replicates of each of the WT and *asrA*-samples after 90 min of tobramycin treatment were collected.

Immunoblot samples were taken corresponding to each sample. The samples were later transferred to –80°C for storage until delivery to the Clinical Proteomics Mass Spectrometry Core Facility at KI/KS.

### Quantitative proteomics

Proteomics analyses were conducted by the Clinical Proteomics Mass Spectrometry Facility, Karolinska Institute/Karolinska University Hospital/Science for Life Laboratory. The samples were lysed and subjected to protein digestion using trypsin followed by multiplex TMT (Tandem Mass Tag) isobaric labelling before loading onto the LC-MS/MS. The multiplexing allowed all the samples in the proteomics set to be analyzed simultaneously. The TMT18-plex was used for the proteomics (18 samples) which included three replicates each of time points 0 and 60 min after protein synthesis shut-off for both VC and *asrA*OE as well as three replicates each of WT and *asrA*-at time point 90 min after addition of tobramycin. The reference genome used for identifying the proteins from unique peptides was that of *P. aeruginosa* PAO1 (NCBI RefSeq: NC_002516.2). The proteomics dataset is available as Supplementary Dataset 1.

### Proteomics data analysis

In the proteomics experiment, a total of 2021 proteins were detected in all samples. The proteins were selected using three criteria: based on reduction in steady-state levels in *asrA*OE, increase in steady-state levels in *asrA*-and degradation over time in *asrA*OE. The data was analyzed for steady-state level changes by calculating the log_2_ fold-change between the 0 min samples of *asrA*OE and VC for each protein and all proteins which show reduced relative levels in *asrA*OE were chosen. Similarly, log_2_ fold-change of WT and *asrA*-at 90 min after tobramycin addition were compared and all proteins with increase in levels in *asrA*-were chosen. Finally, to select candidates that are unstable in *asrA*OE, proteins with the fold-change ≥ 1.05 between 0 min and 60 min in this strain were selected. All criteria were independently applied on the original list of 2021 total proteins detected. To shortlist as many relevant candidates as possible, the selection was not stringent and all the proteins that satisfied these criteria were included without setting more cut-offs. Venn diagram was created using BioVenn (47). The functional analysis of shortlisted candidates was performed using the ontology ‘Protein class’ in the PANTHER 19.0 database on www.pantherdb.org (48) and the unclassified proteins were further assigned functions based on The Pseudomonas Genome Database available on https://www.pseudomonas.com (49).

### Protein purification

Purification of proteins was adapted from Holmberg *et al.* (50) and is in detail described in the Supplementary Methods.

### *In vitro* degradation assay

*In vitro* degradation assays were performed as described previously(8). The reaction was carried out in Lon or AsrA reaction buffer (25 mM Tris-HCl pH 8.0, 100 mM KCl, 10 mM MgCl_2_, 1 mM DTT) using 0.2 µM Lon or AsrA hexamer, respective amounts of substrate as mentioned in the figure legends and an ATP regeneration system (4 mM ATP, 15 mM creatine phosphate, 75 µg/mL creatine kinase). The reaction mix and the ATP regeneration system were prepared separately and pre-warmed to 37°C. The reaction was started by adding the ATP regeneration system to the reaction mix. Samples were taken at indicated time points and quenched by 1 volume of 2× Laemmli SDS loading buffer or 2× Tricine loading buffer (200 mM Tris-HCl, pH6.8, 2% SDS, 40% glycerol, 0.04% Coomassie Brilliant Blue G-250, 2% β-mercaptoethanol) and snap frozen in liquid nitrogen. Samples were heated at 65°C for 10 min and separated by SDS-PAGE (Bio-Rad 4–20% Mini-PROTEAN® TGX or 16.5% Mini-PROTEAN® Tris-Tricine protein gel) stained by ReadyBlue (Sigma-Aldrich), visualized with a Bio-Rad ChemiDoc MP and quantified using Bio-Rad ImageLab 6.0.1. Substrate levels were normalized to the Lon or AsrA levels of the respective time points.

### Plate reader-based growth curve measurements

Overnight cultures of biological replicates were diluted 1:200 in the morning in LB, grown for 2-4 hours, and diluted to an OD_600_ of 0.05. 200 μL of the dilution were added into sterile 96 well transparent plates. LB medium was used as blank in multiple wells. The plate was set up in a Spark microplate reader (Tecan) at 37°C for OD_600_ measurements every 10 min for 24 hours with shaking.

### Spot dilution assays

LB agar plates, containing antibiotics when applicable, were poured and dried for at least 24 hours. Triplicates of respective strains from individual colonies from freshly streaked plates were diluted from overnight cultures 200× into 10 mL LB and grown at 200 rpm at 37°C for 3–4 hours. The cultures were adjusted to an OD_600_ of 0.1 and 10^0^ to 10^-7^ dilutions were made from each culture in a 96-well plate. 1 μL from each dilution of each strain was plated onto an LB plate and incubated at 37°C or 45°C for 24 hours. The plates were imaged using a ChemiDoc under the setting: Blots – Colorimetric and the image was visualized using BioRad Image Lab 6.0.1.

### Swimming motility assay

Swimming motility of *P. aeruginosa* strains was assayed on LB medium plates containing 0.3% agar. After drying the plates for 4h, 1 μL of *P. aeruginosa* strains at OD_600_=0.1was inoculated 3 mm from the surface into the center of the agar. The plates were incubated at room temperature (approximatively 25°C) for 48h. The plates were imaged using a ChemiDoc under the setting: Blots – Colorimetric. Fiji (ImageJ) was used to quantify the motility zones and GraphPad Prism was used to perform statistical analysis.

### Phase contrast microscopy

The strains were grown to exponential phase and samples were collected. 1% final concentration of formaldehyde was used to fix the cells to be stored at 4°C. The cells were transferred to 1% agarose pads on glass slides and covered with cover slips for imaging. A T*i* eclipse inverted research microscope (Nikon) with 100×/1.45 numerical aperture (NA) objective (Nikon) was used to collect phase-contrast images. The images were processed using Fiji (ImageJ).

### Generation of AsrA and QslA antibodies

Polyclonal antibodies from rabbit were produced by Davids Biotechnologie GmbH using purified AsrA and QslA. Purified AsrA and QslA were extracted from a 7 cm prep well Mini-PROTEAN® TGX Stain-Free™ gel (#4568091, Bio-Rad). Polyclonal antiserum was generated in a rabbit by five immunizations. Purified AsrA and QslA were used for affinity purification of the final antibodies.

### Immunoblot analysis

For whole cell extract analysis, 1 mL culture samples were collected at the respective time points and cell pellets were obtained by centrifugation. Cell pellets were resuspended in 200 μL of 1× SDS sample buffer per OD_600_ 1.0, to ensure normalization of the samples. Samples were boiled at 98°C for 10 min and run on an SDS-PAGE using Mini-PROTEAN® TGX Stain-Free^TM^ gels (usually 4-20%, Bio-Rad). The proteins were transferred to nitrocellulose membranes by a semi-dry blotting procedure as per manufacturer guidelines. The protein gels and membranes were imaged using a ChemiDoc MP (Bio-Rad) system before and after the transfer, respectively, to assess equal loading of total protein as well as the quality of the transfer.

Membranes were blocked for 1 hour at room temperature in 10% skim milk powder in TBS (TBS) and protein levels were detected using the anti-Lon (1:10000 dilution; kindly provided by R.T. Sauer) or anti-AsrA (1:100) or anti-QslA (1:67) primary antibodies in 3% skim milk powder in TBS-Tween (TBST). Secondary antibodies, 1:5000 dilutions of anti-rabbit HRP-conjugated (Thermo Fisher Scientific) and SuperSignal Femto West (Thermo Fisher Scientific) were used to detect primary antibody binding. Immunoblots were scanned by chemiluminescence using a ChemiDoc MP (Bio-Rad) system. Relative signal intensities were quantified using Bio-Rad Image Lab 6.0.1.

### Statistical analysis

All quantifications were made from arithmetic mean ± standard deviation of independent replicates. The value of replicate number, sample sizes, and the statistical test used along with p-values are mentioned in the respective figure legends. The application GraphPad Prism Version 10.4.1 was used for the analyses and statistical tests.

## Acknowledgements

We thank members of the Jonas group for helpful discussions and feedback and the Hancock (University of British Columbia) and Ehud Banin (Bar-Ilan University) labs for kindly providing strains. We also thank the Clinical Proteomics Mass Spectrometry Core Facility at KI/KS for excellent service, support and advice. The study was funded by project grants from the Swedish Research Council (2020-03545, 2024-04942 to KJ) and the Swedish Cancer Society (24 3570 to KJ) as well as funding from the Lillian Sagens and Curt Ericssons Research Foundation to UR and from the Strategic Research Area (SFO) program distributed through Stockholm University to KJ. The computations and data handling for the Alphafold3 predictions of QslA-AsrA NTD were enabled by resources provided by the National Academic Infrastructure for Supercomputing in Sweden (NAISS) at Tetralith and Alvis, partially funded by the Swedish Research Council through grant agreement no. 2022-06725, project ID NAISS 2025/22-1088 (to KJ and ENV).

