## Supplementary Methods, Figures S1-S6, Tables S1-S3, Legend of Dataset S1 for "Functional diversification of two Lon homologs enhances stress adaptation in *Pseudomonas aeruginosa*"

The Supplementary Information includes:

Supplementary Methods

Supplementary Figures S1 to S6

Supplementary Tables S1 to S3

Legend of Dataset S1

SI References

### Supplementary Methods

#### Plasmid construction

All plasmids used in this study are listed in Supplementary Table S2.

***asrA overexpression plasmid***

For constructing plasmid pJN105:*asrA*, the *asrA* gene from PAO1 genomic DNA was cloned into pJN105 vector under the control of the L-arabinose–inducible promoter using primers OAK142 and OAK143 (Supplementary Table S3). The gene was inserted between restrictions sites of NheI and PstI through restriction digestion and Gibson assembly of the fragments. The DNA assembly was subsequently transformed into *E. coli* DH5α competent cells and transformants selected on LB plates containing gentamicin. The plasmid was isolated and sequence-verified by Sanger sequencing (Eurofins Genomics) with primers OAK145 and OAK146 (Supplementary Table S3). The pJN105 empty vector (VC) and cloned overexpression plasmid were inserted through electroporation into electrocompetent *P. aeruginosa* cells of WT background to generate the vector control or overexpression strain. The successful transformants were selected on LB plates containing gentamicin after overnight growth at 37°C.

***Protein expression plasmids***

Plasmids for protein expression were generated by amplifying the corresponding gene from the genomic DNA of strain PAO1 using primer pairs as listed in Supplementary Table S3 and subsequent cloning of the insert into a pSUMO-YHRC vector backbone by Gibson assembly. The plasmid was amplified as two fragments from the original using primer pairs OMJF34/OMJF36 and OMJF37/OMJF38 (Supplementary Table S3) disrupting the kanamycin resistance cassette. The two plasmid fragments and the amplified gene were then joined using Gibson assembly and the assembly product transformed into chemically competent DH5α cells, followed by selection on LB plates containing kanamycin. The plasmid was then extracted and sequence-verified by Sanger sequencing (Eurofins Genomics) with primers OAK31 and OAK32. The primer pairs used for cloning of each gene are as listed in Supplementary Table S3.

#### Protein purification

Purification of proteins was adapted from Holmberg *et al.* (1). BL21-SI/pCodonPlus cells were transformed using the plasmids mentioned in the Supplementary Table S2 by electroporation and selected on LBON agar plates supplemented with kanamycin (Kan) and chloramphenicol (Chlor). Pre-cultures (LBON or 2xYTON + Kan + Chlor) were inoculated with about 20 colonies and cultivated at 30°C overnight. One liter of 2xYTON + Kan + Chlor was inoculated by 1:100 dilution of the pre-culture and cultured until an approximate $OD_{600}$ of 1.0. The expression was started by addition of 0.5 mM IPTG and 0.3 M NaCl (final concentrations). Incubation continued at 30°C for up to 4 hours and cells were harvested subsequently by centrifugation (6 800×g, 4°C, 10 min) and cell pellets stored at −80°C.

For purification, pellets were resuspended in lysis buffer (40 mM HEPES-KOH pH 7.5, 500 mM NaCl or KCl, 10% glycerol) supplemented with 1 mM PMSF, 1 mg/mL Lysozyme and 3 µL Benzonase/10 mL and topped up to 30 mL total volume. Cells were then lysed by 2–

3 passes through an EmulsiFlex-C3 high-pressure homogenizer and peak pressure was kept between 25 000 and 30 000 psi. Lysate was cleared by centrifugation at 32 500×g at 4°C for 1.5 h. Tagged proteins were bound to 3 mL (=1.5 mL bed volume) HisPur Cobalt Resin (Thermo Fisher Scientific, 89965) per liter culture, pre-equilibrated with lysis buffer, for 30 min at 4°C with shaking. After washing 5 times with approximately 50 mL of lysis buffer containing 10 mM imidazole, bound proteins were eluted using elution buffer with 250 mM imidazole. Fractions with protein concentrations ≥ 0.2 mg/mL as measured using a Nanodrop, were pooled. For removal of imidazole, the pooled protein was transferred into dialysis membranes with cut-off of approximatively one-third of the respective protein's molecular weight. The dialysis was performed against storage buffer (40 mM HEPES-KOH pH 7.5, 500 mM NaCl or KCl, 10% glycerol) overnight at 4°C with constant stirring. For proteins that require removal of the 6×His-SUMO tag, 4 µg/mL Ulp1-6×His protease was added to the fractions during the dialysis. Tag depletion was achieved by binding to 3 mL (=1.5 mL bed volume) pre-equilibrated HisPur Cobalt Resin from (Thermo Fisher Scientific, 89965). Flow through and/or wash fractions (imidazole gradient) containing purified protein was collected. Lon and AsrA were further purified by anion exchange with a HiTrap Q HP 1 mL column (Cytiva, purchased on VWR: 29-0513-25) in a ÄKTA FPLC system at a flow rate of 1 mL/min. The column was washed and the proteins were bound by using buffer A (50 mM Tris/Cl pH 8.0, 50 mM KCl, 10% glycerol) and buffer B (50 mM Tris/Cl pH 8.0, 1 M KCl, 10% glycerol). Lon and AsrA were eluted with 40% and 20% buffer B, respectively. Protein fractions were subsequently analyzed by SDS-PAGE (Bio-Rad 4–20% Mini-PROTEAN® TGX protein gel), stained by ReadyBlue (Sigma-Aldrich), visualized with a Bio-Rad ChemiDoc MP and quantified using Bio-Rad ImageLab 6.0.1. Protein concentrations were calculated comparing quantified SDS-PAGE band intensities against a BSA standard curve and/or by NanoDrop (absorbance at 280 nm). When necessary, proteins were concentrated using a Pall Advanced Centrifugal Device or an Amicon Ultra Filter with a pore size around one-third of the respective protein molecular weight. Before storage, 1 mM DTT and 0.1 mM EDTA was added to the protein. Proteins were snap-frozen in liquid nitrogen and stored at −80°C.

### Supplementary Figures

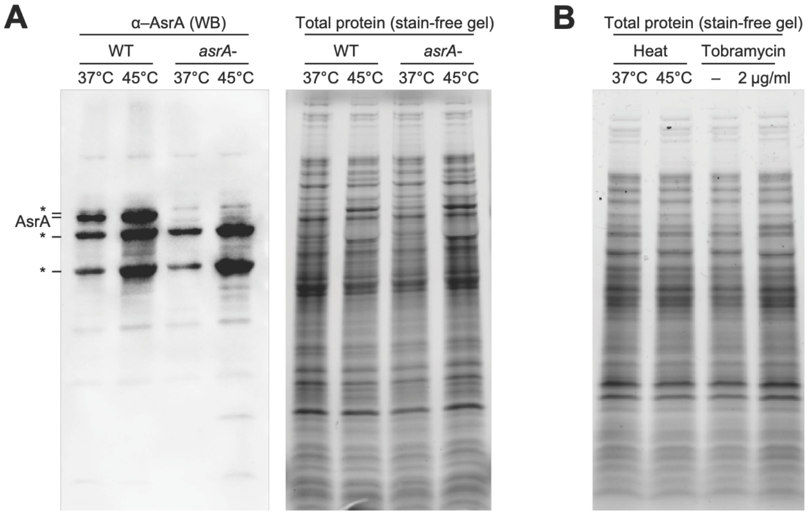


**Figure S1. Additional controls demonstrating the specificity of the AsrA antibody and protein loading.**

**A)** *Left:* Representative western blot (WB) showing AsrA antibody signals in the *P. aeruginosa* PAO1 WT and *asrA-* before and after 30 min of heat stress. *Right:* Corresponding stain-free gel showing total protein loading. **B)** Representative stain-free gel corresponding to the western blot shown in **Fig. 1B**, with protein samples from *P. aeruginosa* PAO1 WT cells before and after heat stress and 2 µg/mL tobramycin treatment for 30 min.

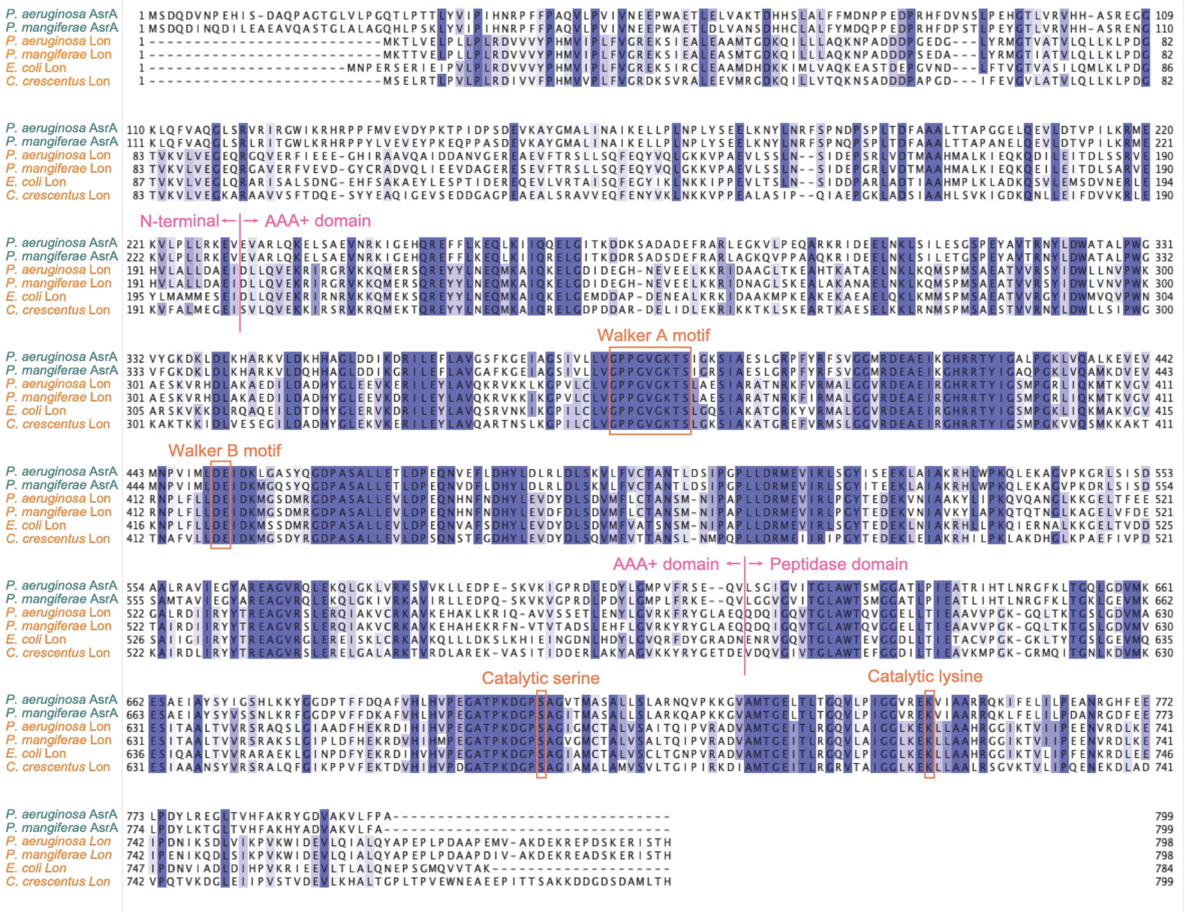


**Figure S2. AsrA alignment with Lon protease orthologs.**

Alignment of *Pseudomonas aeruginosa* AsrA (PA0779) with *Pseudomonas mangiferae* AsrA, *Pseudomonas aeruginosa* Lon (PA1803), *Pseudomonas mangiferae* Lon, *Escherichia coli* Lon, and *Caulobacter crescentus* Lon. The overall conservation between the analyzed sequences is displayed in blue (dark blue: complete conservation and light blue: partial conservation). The important functional amino acids are highlighted in red boxes like the Walker AB motifs in the AAA+ domain and the serine-lysine catalytic dyad in the protease domain. The alignment was generated using Clustal Omega (2) and visualized using Jalview 2.11.5.1.

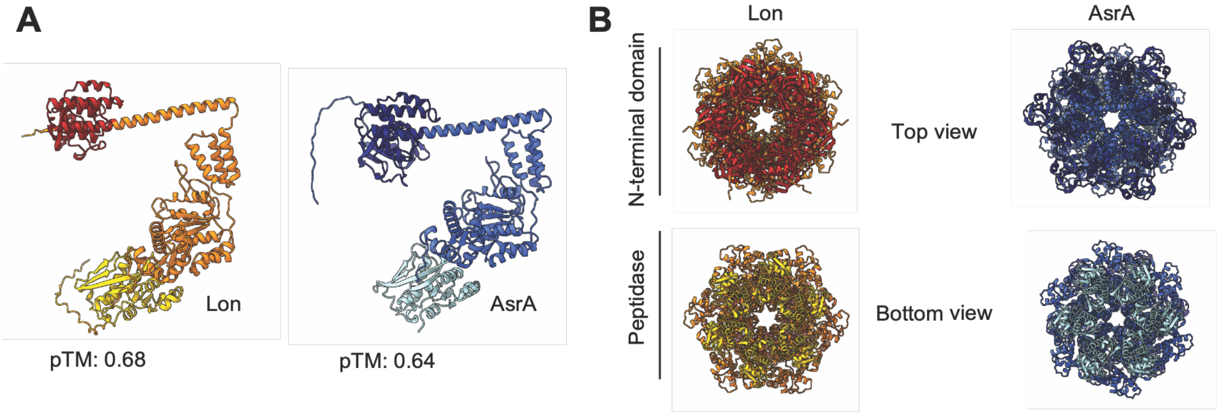


**Figure S3. Lon and AsrA predicted 3D structures.**

**A)** AlphaFold 3 predicted structure of monomeric Lon and AsrA. N-terminal domain is displayed in dark red (Lon) and dark blue (AsrA), AAA+ unfoldase domain in orange (Lon) and blue (AsrA), peptidase domain in yellow (Lon) and light blue (AsrA). **B)** Top and bottom views of predicted hexameric Lon and AsrA shown in **Fig. 1D** (pTM: predicted Template Modeling score).

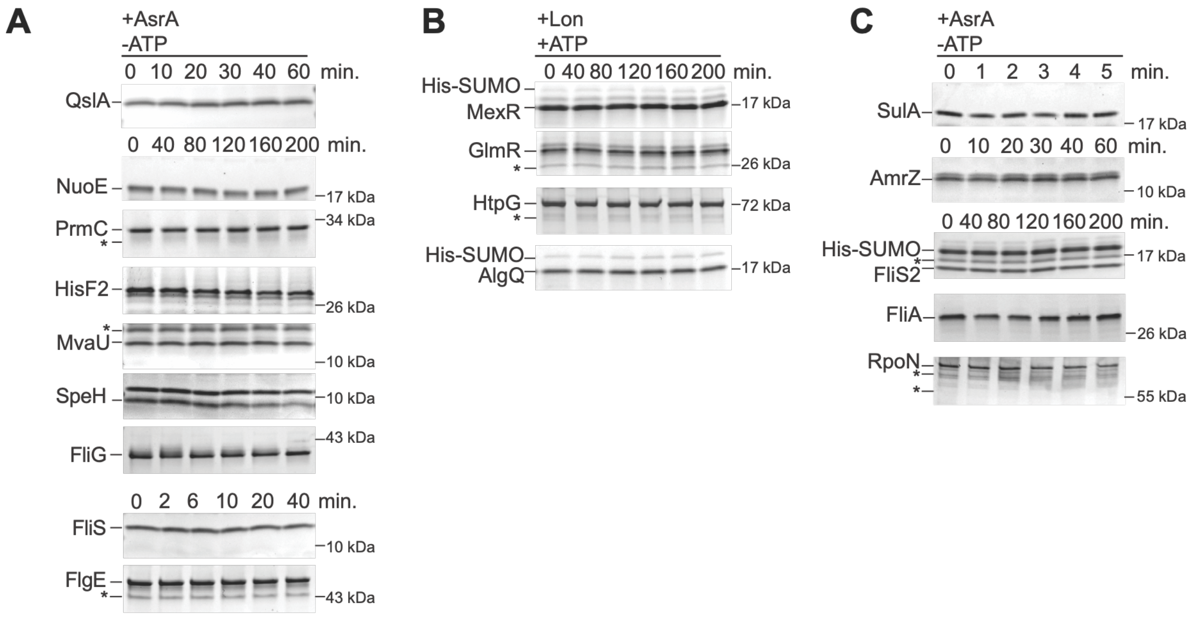


**Figure S4. Additional *in vitro* degradation assays of AsrA substrates.**

**A)** *In vitro* degradation assays of the novel AsrA substrates displayed in **Fig. 3B** in the absence of ATP and ATP regeneration system. One representative SDS-PAGE gel of each assay is shown. **B)** *In vitro* degradation assays of MexR, GlmR, HtpG and AlgQ in the presence of Lon, ATP, and ATP regeneration system. 0.2 µM of Lon hexamer and 2 µM of substrates (except AlgQ that was added at a concentration of 3 µM) were used in all assays. **C)** *In vitro* degradation assays of the Lon and AsrA substrates displayed in **Fig. 3D** in the absence of ATP and ATP regeneration system.

| Protein | Function | No. satisfied criteria | Prev. validated Lon substrate |
|---|---|---|---|
| FliA | RNA polymerase sigma factor FliA | 2 | Yes |
| RpoN | RNA polymerase sigma-54 factor | 1 | Yes |
| AmrZ | Transcription factor AmrZ | ND | Yes |
| FliS2 | Hypothetical protein FliS-like* | / | Yes |
| SulA | Cell division inhibitor SulA | ND | Yes |

*from *Pseudomonas aeruginosa* clone C 8277

**Figure S5. Validated Lon substrates not classified as putative AsrA substrates.**

Table of previously validated Lon substrates analyzed in **Fig. 3D-E** that did not belong to the 124 putative AsrA substrates identified in the proteomics analysis. Numbers of satisfied criteria in the proteomics experiment are indicated along with the primary function in the cell (ND: not detected in this proteomics experiment; /: absent in *Pseudomonas aeruginosa* PAO1 strain).

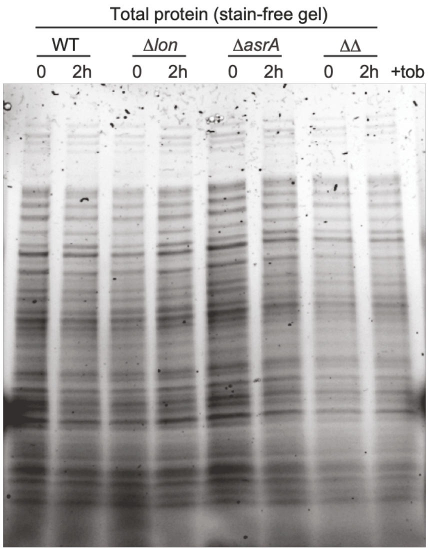


**Figure S6. Representative stain-free gel corresponding to the western blot shown in Fig. 4F**. The gel shows protein samples from *P. aeruginosa* PAO1 WT cells and the Δ*lon*, Δ*asrA* and Δ*asrA* Δ*lon* (ΔΔ) strains before and after 2 µg/mL tobramycin treatment for 2 hours.

### Supplementary Tables

**Table S1 – Strains used in this study.**

| ***Escherichia coli* strains** | | | | |
|---|---|---|---|---|
| **Name** | **Genotype** | **Description** | **Marker** | **Reference** |
| DH5α | General cloning strain | - | - | Invitrogen |
| BL21-SI/ pCodonPlus | Salt-inducible BL21(DE3) strain for protein expression | - | chlor$^{R}$ | Provided by Claes Andréasson, Stockholm University, Sweden |
| ***Pseudomonas aeruginosa* strains** | | | | |
| **Name** | **Genotype** | **Description** | **Marker** | **Reference** |
| H103 | PAO1 WT | WT | - | Provided by Robert Hancock, University of British Columbia, Canada |
| H1105 | PAO1 mini-Tn*5*–*luxCDABE*::*lon*; 74_D9, (PA1803) | *lon*- mutant | tet$^{R}$ | (3) |
| lux_15_F1 | PAO1 mini-Tn5-*luxCDABE*::*asrA*; 74_D9, (PA0779) | *asrA- mutant* | tet$^{R}$ | (3) |
| KJ1230 | PAO1 WT + pJN105 | WT transformed with empty vector pJN105 | gent$^{R}$ | (4) |
| KJ1235 | PAO1 WT + pAK023 | WT transformed with pJN105 containing *asrA*; *asrA* overexpression strain | gent$^{R}$ | This study |
| | PAO1 WT | WT | - | Provided by Ehud Banin, Bar-Ilam University, Israël |
| | PAO1 Δ*lon* (PA1803) | *lon* deletion | - | (5) |
| | PAO1 Δ*asrA* (PA0779) | *asrA* deletion | - | (5) |

| | | | | |
|---|---|---|---|---|
| | PAO1 Δ*asrA* Δ*lon* | *asrA* and *lon* deletion | - | (5) |

**Table S2 – Plasmids used in this study.**

<table>
<tr><th>Name</th><th>Description</th><th>Marker</th><th>Reference</th></tr>
<tr><td>pJN105</td><td>Broad-host range vector with L-arabinose-inducible araBAD promoter; pBBR1ori</td><td>gent$^{R}$</td><td>(6)</td></tr>
<tr><td>pAK023</td><td>pJN105 containing asrA</td><td>gent$^{R}$</td><td>This study</td></tr>
<tr><td>pSUMO-YHRC</td><td>Plasmid for protein expression using $P_{T7}$ with an N-terminal 6xHis-SUMO tag; RRID: Addgene_54336</td><td>kan$^{R}$</td><td>(1)</td></tr>
<tr><td>pAK010</td><td>pSUMO-YHRC containing 6xHis-SUMO-lon</td><td>kan$^{R}$</td><td rowspan="10">(4)</td></tr>
<tr><td>pAK011</td><td>pSUMO-YHRC containing 6xHis-SUMO-sulA</td><td>kan$^{R}$</td></tr>
<tr><td>pAK012</td><td>pSUMO-YHRC containing 6xHis-SUMO-fliG</td><td>kan$^{R}$</td></tr>
<tr><td>pAK013</td><td>pSUMO-YHRC containing 6xHis-SUMO-flgE</td><td>kan$^{R}$</td></tr>
<tr><td>pAK014</td><td>pSUMO-YHRC containing 6xHis-SUMO-fliA</td><td>kan$^{R}$</td></tr>
<tr><td>pAK015</td><td>pSUMO-YHRC containing 6xHis-SUMO-rpoN</td><td>kan$^{R}$</td></tr>
<tr><td>pAK016</td><td>pSUMO-YHRC containing 6xHis-SUMO-amrZ</td><td>kan$^{R}$</td></tr>
<tr><td>pAK017</td><td>pSUMO-YHRC containing 6xHis-SUMO-fliS</td><td>kan$^{R}$</td></tr>
<tr><td>pAK018</td><td>pSUMO-YHRC containing 6xHis-SUMO-fliS2</td><td>kan$^{R}$</td></tr>
<tr><td>pAK020</td><td>pSUMO-YHRC containing 6xHis-SUMO-speH</td><td>kan$^{R}$</td></tr>
<tr><td>pAK024</td><td>pSUMO-YHRC containing 6xHis-SUMO-asrA</td><td>kan$^{R}$</td><td>This study</td></tr>
<tr><td>pAK025</td><td>pSUMO-YHRC containing 6xHis-SUMO-hisF2</td><td>kan$^{R}$</td><td>This study</td></tr>
<tr><td>pAK026</td><td>pSUMO-YHRC containing 6xHis-SUMO-mexR</td><td>kan$^{R}$</td><td>This study</td></tr>
<tr><td>pAK027</td><td>pSUMO-YHRC containing 6xHis-SUMO-qslA</td><td>kan$^{R}$</td><td>This study</td></tr>
<tr><td>pAK034</td><td>pSUMO-YHRC containing 6xHis-SUMO-mvaU</td><td>kan$^{R}$</td><td>This study</td></tr>
<tr><td>pML192</td><td>pSUMO-YHRC containing 6xHis-SUMO-prmC</td><td>kan$^{R}$</td><td>This study</td></tr>
</table>

| | | | |
|---|---|---|---|
| pML196 | pSUMO-YHRC containing *6xHis-SUMO-glmR* | $kan^R$ | This study |
| pML198 | pSUMO-YHRC containing *6xHis-SUMO-htpG* | $kan^R$ | This study |
| pML199 | pSUMO-YHRC containing *6xHis-SUMO-algQ* | $kan^R$ | This study |
| pML201 | pSUMO-YHRC containing *6xHis-SUMO-nuoE* | $kan^R$ | This study |

**Table S3 – Oligonucleotides used in this study.**

| Name | Sequence (5'-3') | Description | Reference |
|---|---|---|---|
| OAK031 | TAATACGACTCACTATAGGG | T7 promoter forward | common primer |
| OAK032 | GCTAGTTATTGCTCAGCGG | T7 terminator reverse | common primer |
| OAK033 | AGGGTTTTCCCAGTCACGACGTT | M13 reverse | common primer |
| OAK034 | GAGCGGATAACAATTTCACACAG | M13 forward | common primer |
| OAK037 | CACAGAGAACAGATTGGTGGATGAAAACACTCGTCGAATTGC | *lon* insert for pSUMO-YHRC forward | |
| OAK038 | GACGGAGCTCGAATTCGGATCCTAATGCGTGCTAATTCGCTC | *lon* insert for pSUMO-YHRC reverse | |
| OAK039 | CACAGAGAACAGATTGGTGGATGCAGACCTCCCACTCGCTG | *sulA* insert for pSUMO-YHRC forward | |
| OAK040 | GACGGAGCTCGAATTCGGATCTCAACCCAGACGAATATTCAG | *sulA* insert for pSUMO-YHRC reverse | |
| OAK091 | CACAGAGAACAGATTGGTGGATGAGTGAGAATCGTCTCGCCG | *fliG* insert for pSUMO-YHRC forward | |
| OAK092 | GACGGAGCTCGAATTCGGATCTCAGATCATCTCCTCGCCACCCTTG | *fliG* insert for pSUMO-YHRC reverse | (4) |
| OAK134 | CACAGAGAACAGATTGGTGGATGAGTTTCAACATCGGCCTGAGCGGCATCCAGGC | *flgE* insert for pSUMO-YHRC forward | |
| OAK135 | GAGCTCGAATTCGGATCTCAGCGCAGGTTGATGATGGTCTGGGTCACCGCATCCTCGGTC | *flgE* insert for pSUMO-YHRC reverse | |
| OAK136 | CACAGAGAACAGATTGGTGGATGACAGCGGCCTCTGGAGTGCGTATGTATAGC | *fliA* insert for pSUMO-YHRC forward | |
| OAK137 | GAGCTCGAATTCGGATCTCAGGCCGACCGCCAATCGGCCAGGCGCGCGCAAAC | *fliA* insert for pSUMO-YHRC reverse | |
| OAK138 | CACAGAGAACAGATTGGTGGATGAAACCATCGCTAGTCCTCAAGATGGCCAGC | *rpoN* insert for pSUMO-YHRC forward | |

| | | | |
|---|---|---|---|
| OAK139 | GAGCTCGAATTCGGATCTCACACCAGTCGC TTGCGCTCGCTCGAAGGGGC | *rpoN* insert for pSUMO-YHRC reverse | |
| OAK140 | CACAGAGAACAGATTGGTGGGATGCGCCCA CTGAAACAGGCAACTCCTACCTAC | *amrZ* insert for pSUMO-YHRC forward | |
| OAK141 | GAGCTCGAATTCGGATCTCAGGCCTGGGCC AGCTCCGCATCGTGTGCGATC | *amrZ* insert for pSUMO-YHRC reverse | |
| OAK175 | CACAGAGAACAGATTGGTGGGATGAACGCA ATGGCAGCCATGCGGCAATAC | *fliS* insert for pSUMO-YHRC forward | |
| OAK176 | GAGCTCGAATTCGGATCTCAGGGGGCAATC GCATCCCAACCGGATTTGATG | *fliS* insert for pSUMO-YHRC reverse | |
| OAK177 | CACAGAGAACAGATTGGTGGGATGTCGCGT CCTATCGATACCTACCGGCAG | *fliS2* insert for pSUMO-YHRC forward | |
| OAK178 | GGAGCTCGAATTCGGATCTTAGCGCCGTTC GCTGTCTTCGCCCTGCGCTTCG | *fliS2* insert for pSUMO-YHRC reverse | |
| OAK109 | CACAGAGAACAGATTGGTGGGATGGCAATT CAACCGTTGCGACTCGATCCG | *speH* insert for pSUMO-YHRC forward | |
| OAK110 | GACGGAGCTCGAATTCGGATCTCAGGCCAC CCCCGTGCCCTGGCCCAGCGC | *speH* insert for pSUMO-YHRC reverse | |
| OAK128 | CACAGAGAACAGATTGGTGGGATGAGCGAC CAGGACGTTAATC | *asrA* insert for pSUMO-YHRC forward | This study |
| OAK129 | GACGGAGCTCGAATTCGGATCTCAGGCCGG GAACAGCACCTTG | *asrA* insert for pSUMO-YHRC reverse | This study |
| OAK132 | GCTGGCCCTGTTCTTCATGG | Sequencing *asrA* insert for pSUMO-YHRC forward | This study |
| OAK133 | TCCCTGGGACCGATCTTCAC | Sequencing *asrA* insert for pSUMO-YHRC reverse | This study |
| OAK142 | CGCAACTCTCTACTGTTTCTCCATACCCGT TTTTTTGGGCTAGCATGAGCGACCAGGACG TTAATCCCGAACAC | pJN105 NheI *asrA*OE cloning forward | This study |

| | | | |
|---|---|---|---|
| OAK143 | CGCGTAATACGACTCACTATAGGGCGAATT GGAGCTCTCAGGCCGGGAACAGCACCTTGG CCACGTCGCCGTAG | pJN105 SacI *asrA*OE cloning reverse | This study |
| OAK145 | GTAACAAAGCGGGACCAAAG | Sequencing *asrA*OE insert in pJN105 forward | This study |
| OAK146 | CAACTGTTGGGAAGGGCGATC | Sequencing *asrA*OE insert in pJN105 reverse | This study |
| OAK205 | GATATTATTGAGGCTCACAGAGAACAGATT GGTGGGATGGTCCGGAGGCGCGTTATCCCA TGCTTGCTGCTCAAG | *hisF2* insert for pSUMO-YHRC forward | This study |
| OAK206 | CTTGTCGACGGAGCTCGAATTCGGATCCTA ACCGACGTCGAGCTTGTTGACATCCGGATA ACTAATCAGTACCGC | *hisF2* insert for pSUMO-YHRC reverse | This study |
| OAK199 | GATATTATTGAGGCTCACAGAGAACAGATT GGTGGGATGAACTACCCCGTGAATCCCGAC CTGATGCCCGCGCTGATG | *mexR* insert for pSUMO-YHRC forward | This study |
| OAK200 | CTTGTCGACGGAGCTCGAATTCGGATCTTA AATATCCTCAAGCGGTTGCGCGGCCAGGCA CTGGTCGAG | *mexR* insert for pSUMO-YHRC reverse | This study |
| OAK209 | GATATTATTGAGGCTCACAGAGAACAGATT GGTGGGGTGACTCTCAGAAACGGAGTACCC AGCATGACGAAGGATG | *qslA* insert for pSUMO-YHRC forward | This study |
| OAK210 | CTTGTCGACGGAGCTCGAATTCGGATCTCA ACCGGAACGTCGAGCGGCTACCAGGCGCTG CTGCAGGCGCGTCAG | *qslA* insert for pSUMO-YHRC reverse | This study |
| OAK219 | GATATTATTGAGGCTCACAGAGAACAGATT GGTGGGATGTCCAAACTTGCCGAGTTCCGC GAGGCAGAGCGCAAAC | *mvaU* insert for pSUMO-YHRC forward | This study |
| OAK220 | CTTGTCGACGGAGCTCGAATTCGGATCTTA GCGTTGCAGCCAGGATTCGACGGTTTCGGA ACCGTACTGTTCTTTC | *mvaU* insert for pSUMO-YHRC reverse | This study |
| OML146 | TAACGATATTATTGAGGCTCACAGAGAACA GATTGGTGGGATGACCACTATCTGTACCCT TCTCAAGGATTCCCA | *prmC* insert for pSUMO-YHRC forward | This study |
| OML147 | GTGCGGCCGCAAGCTTGTCGACGGAGCTCG AATTCGGATCTCAGCATGCCCATTGTCCGA | *prmC* insert for pSUMO-YHRC reverse | This study |
| OML150 | TAACGATATTATTGAGGCTCACAGAGAACA GATTGGTGGGGTGATCTCGAAACGTAATAC GCCGCAACGTCGCC | *glmR* insert for pSUMO-YHRC forward | This study |

| | | | |
|---|---|---|---|
| OML151 | GTGCGGCCGCAAGCTTGTCGACGGAGCTCG AATTCGGATCTCAGGCCTCGACCGGGGTGC | *glmR* insert for pSUMO-YHRC reverse | This study |
| OML154 | TAACGATATTATTGAGGCTCACAGAGAACA GATTGGTGGGATGAGCGTGGAAACCCAAAA AGAGACACTG | *htpG* insert for pSUMO-YHRC forward | This study |
| OML155 | GTGCGGCCGCAAGCTTGTCGACGGAGCTCG AATTCGGATCTTAAGCGGACAGCTCCACCA | *htpG* insert for pSUMO-YHRC reverse | This study |
| OML156 | TTCCACGGCGAAGAGGCCGACAAGCC | Sequencing *htpG* insert in pSUMO-YHRC forward | This study |
| OML157 | CTCCTTGTTGCCGAAGTCTTCCGCC | Sequencing *htpG* insert in pSUMO-YHRC reverse | This study |
| OML160 | TAACGATATTATTGAGGCTCACAGAGAACA GATTGGTGGGATGCTCGAAAGCTGCCGTAA TGCCCAA | *algQ* insert for pSUMO-YHRC forward | This study |
| OML161 | GTGCGGCCGCAAGCTTGTCGACGGAGCTCG AATTCGGATCTCAGACCGGTACTGCCGAAC | *algQ* insert for pSUMO-YHRC reverse | This study |
| OML168 | TAACGATATTATTGAGGCTCACAGAGAACA GATTGGTGGGATGAGCCAGAGCAACCTGAT TCAGACCGAC | *nuoE* insert for pSUMO-YHRC forward | This study |
| OML169 | GTGCGGCCGCAAGCTTGTCGACGGAGCTCG AATTCGGATCTCATACGTAGGCCTCCAGCA | *nuoE* insert for pSUMO-YHRC reverse | This study |
| OMJF34 | CCCACCAATCTGTTCTCTGTG | Cloning pSUMO-YHRC fragment 1 forward | (7) |
| OMJF36 | CATGCATCATCAGGAGTACGG | Cloning pSUMO-YHRC fragment 1 reverse | |
| OMJF37 | GATCCGAATTCGAGCTCC | Cloning pSUMO-YHRC fragment 2 forward | |
| OMJF38 | GAATTTATGCCTCTTCCGACC | Cloning pSUMO-YHRC fragment 2 reverse | |

### Supplementary Dataset Legends

**Supplementary Dataset 1 –** Quantitative proteomics data, related to Figures 2 and 4.
